# Analysis of Classic Tomato Mutants Reveals Influence of Leaf Vein Density on Fruit BRIX

**DOI:** 10.1101/2021.03.01.433399

**Authors:** Zizhang Cheng, S.D. Rowland, Karo Czarnecki, Kristina Zumstein, Hokuto Nakayama, Neelima R. Sinha

## Abstract

Tomato *bipinnate* (*bip*) is a classic leaf mutant, with highly increased leaf complexity resulting from the loss of function of a *BEL-LIKE HOMEODAMAIN* (*BELL*) gene. Here, we analyzed several *bip* mutants and their isogenic wildtype backgrounds for a suite of leaf morphology traits, ranging from leaf complexity, leaflet shape and size, to leaf vascular density to investigate how changes in leaf morphology influence fruit traits. Our analyses showed an unexpected relationship between leaf vein density and fruit sugar levels, where leaf vein density was negatively correlated with fruit BRIX. RNA-Seq analysis suggested variation in *Glucose-6-phosphate translocator2* (*GPT2*) gene expression caused correlated changes in leaf vein density and BRIX when *bip* mutant and wildtype were compared, suggesting that the correlation between leaf vein density and fruit sugar may result from the genes regulating leaf vein development that are also involved in regulating leaf sugar biosynthesis. Our results provide a resource for further exploration of the genetic basis for the complex relationship between fruit quality and leaf traits in natural populations.

## INTRODUCTION

Tomato (*Solanum lycopersicum*) is one of the highest-value and most extensively used vegetable crops worldwide. However, to meet increasing demand, modern tomato cultivars have been selected for qualities such as size and firmness instead of taste (Beullens et al., 2008; Wang and Seymour, 2017). Consequently, most of modern commercial varieties have lost their flavor and are often tasteless (Klee and Tieman, 2013; Tieman et al., 2017).

Flavor of fruit is the sum of interactions between taste and aroma, whereas sugars and acids are the two of primarily components to activate taste receptors and aroma components such as volatile compounds activate olfactory receptors (Malundo et al., 1996; Baldwin et al., 2008; Tieman et al., 2012). Though the relative contribution of taste and aroma to fruit flavor has not been clearly defined (Malundo et al., 1996), plenty of studies have shown the importance of sugars and acids in determining fresh fruit flavor (Malundo et al., 1996; Baldwin et al., 2008; Beckles et al., 2012). For tomato, the levels of sugars and acids not only contribute to tomato taste (sweetness and sourness), but also are major factors affecting tomato overall flavor intensity (Allen Stevens, 1979; Jones and Scott, 1984), and increasing sugar content of the fruit will enhance tomato flavor (Malundo et al., 1996; Tieman et al., 2017). Recent studies have shown that fruit sugar accumulation in modern tomato is two to three-fold less than that in wild species (Beckles et al., 2012), which can account for the decline in flavor quality of tomato fruit.

Fruits are the primary photosynthetic sinks and over 80% of sugars in the fruit are produced in the leaf through photosynthesis and subsequently translocated through the phloem (Cocaliadis et al., 2014). Therefore, factors involved in regulating leaf photosynthesis, as well as sugar biosynthesis and sugar transport would have an effect on sugar levels in fruit. Leaves are the principle site of plant photosynthesis and leaf traits (e.g. shape and size) directly impact the efficiency of light capture and photosynthetic carbon fixation (Smith et al., 1997; Sarlikioti et al., 2011). Thus, changes in leaf traits could have an effect on fruit yield and quality. Studies evaluating the influence of leaf area on tomato yield have shown high leaf area index (LAI) can lead to an increase in tomato yield as a result of better light interception (Heuvelink et al., 2005). Recently, leaf shape was shown to be strongly correlated with fruit sugar levels in tomato, with rounder and more circular leaves having higher sugar content in their fruit (Chitwood et al., 2013; Rowland et al., 2019). These studies focused on the influence of leaf morphology on fruit sugar level, and revealed important correlations between leaf shape and fruit sugar accumulation. However, how leaf shape contributes to sugar accumulation in fruit is not yet known (Malundo et al., 1996; Chitwood et al., 2013). The impact of leaf traits such as leaf complexity and leaf veins (as the direct conduit of sugar transport) on fruit sugar levels is currently uninvestigated.

Tomato *bipinnate* (*bip*) is a classic leaf mutant, with highly increased leaf complexity (Fig. 1A) resulting from the loss of function of a *BEL-LIKE HOMEODAMAIN* (*BELL*) gene (Kimura et al., 2008) called *BIPINNATE* (*BIP*, Solyc02g089940) (Fig. 1B), and provides ideal material to investigate the influence of leaf traits on fruit sugars. To investigate the links between leaf complexity and fruit sugar accumulation, we performed an analysis of leaflet shape, leaf complexity, vein density, yield, and fruit BRIX on *bip* mutants (*bip2* and *bip0663*) and their isogenic backgrounds (M82 and Lukullus, respectively). Results suggested that leaf vein density, and not leaf complexity, was highly correlated to fruit BRIX in *bip* mutants. RNA-Seq analysis of gene expression in the vegetative apices and leaves of *bip* mutants and wildtypes shows an association between the expression of genes regulating leaf vein development and carbohydrate metabolism and transport, suggesting that alteration in leaf vein development would also modulate processing of carbohydrate metabolism and transport which, in turn, affects the accumulation of fruit sugar. Our analysis offers insight into how leaf morphology may influence fruit sugar and provides a new direction to improve tomato fruit sugar content.

**Fig.1.**
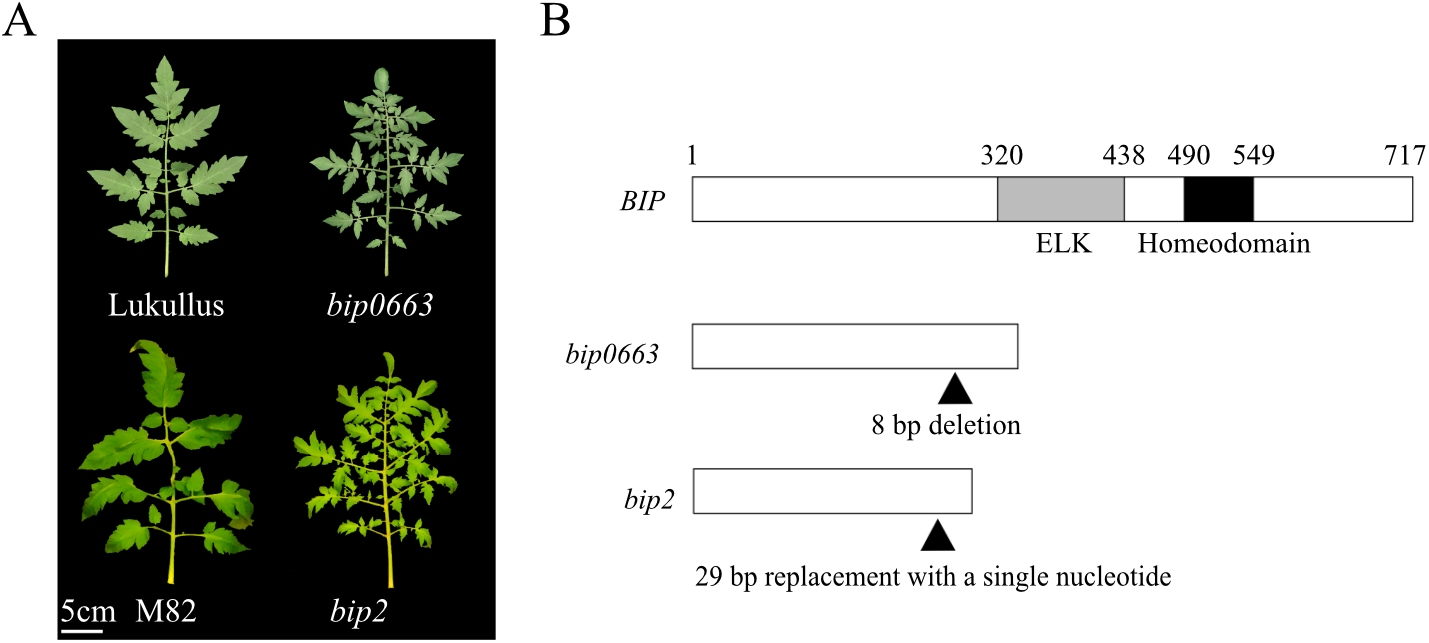
Leaf morphology of *bip* mutants and diagram of *bip* mutations. A, Leaves of *bip* mutants and their corresponding wildtype cultivars; B, Wildtype *BIP* gene with ELK and Homeodomain, represented by gray and black boxes, respectively. Diagram of BIP protein of *bip0663* and *bip2*.

## RESULTS

### Leaf Complexity and Fruit BRIX of *bip* Mutants and Wildtype

To investigate whether changes of leaf complexity caused by mutations at the *BIP* gene have an effect on fruit BRIX, we measured leaf complexity and fruit BRIX of *bip0663* mutants and the corresponding isogenic cultivar, Lukullus. Leaf complexity and fruit BRIX of mutants *bip0663* were both significantly increased in comparison to that of Lukullus (Fig.2A and 2B). However, another *bip* mutant, *bip2*, had significantly increased leaf complexity (Fig.2A), but fruit BRIX similar to its isogenic cultivar, M82 (Fig.2B). These results indicate changes in leaf complexity are not directly correlated fruit BRIX. Additionally, fruit BRIX of both *bip0663* and Lukullus were respectively higher than that of *bip2* and M82. Fruit yield was also measured, since yield has an impact on fruit BRIX (with a general negative correlation between yield and BRIX) (Caliman et al., 2007). We saw fruit yield in *bip0663* similar to Lukullus, while *bip2* fruit yield was lower than M82, suggesting that the difference between *bip0663* and Lukullus BRIX was not directly related to variation in yield. Similar to fruit BRIX, fruit yield of *bip0663* and Lukullus was also significantly higher than that of *bip2* and M82. This result was confirmed by sampling across two seasons (2016 and 2017) in the field and in greenhouse experiments (**Figure S2**).

**Fig.2.**
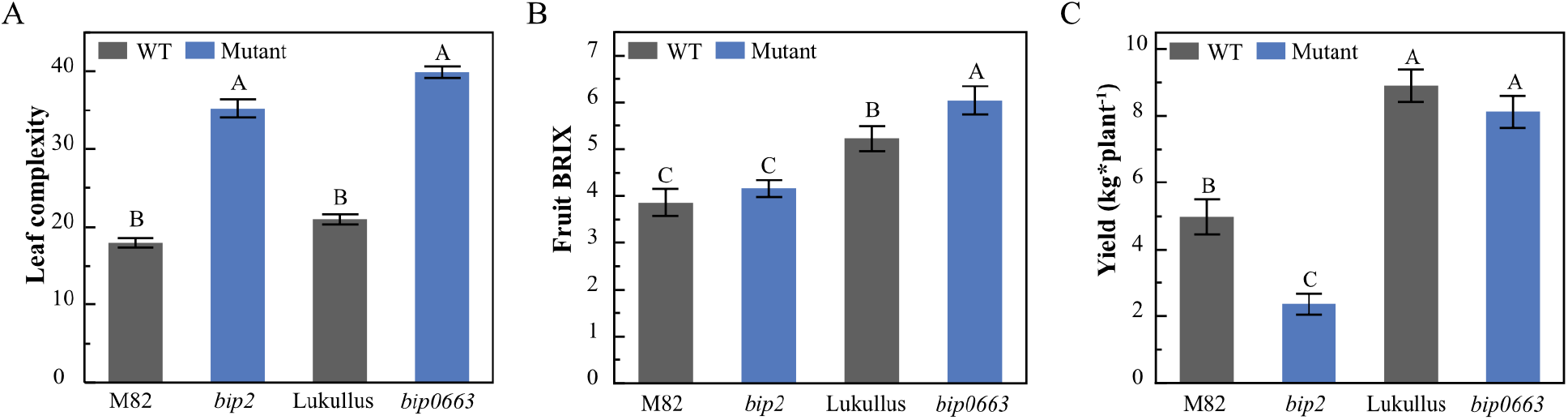
Leaf complexity, fruit BRIX, and yield of *bip* mutants and their corresponding isogenic cultivars. A, Leaf complexity is denoted by the number of all leaflets present on the leaf; B and C, Fruit BRIX (B) and fruit yield (C) of *bip* mutants and their corresponding isogenic cultivars.

### Influence of Leaf Traits on fruit BRIX between *bip0663* and Lukullus

To validate whether changes in leaf traits contributed to the changes in fruit BRIX between *bip0663* and Lukullus, we performed a grafting assay where the Lukullus and *bip0663* separately functioned as reciprocal scion and rootstock. During flowering and fruiting stage, leaves on scion and flowers and fruits on rootstock were completely removed. Thus, the flowering and fruiting of scion were driven by photosynthate transfer only from the leaves on the rootstock portion. In the control group, the Lukullus and *bip0663* were self-grafted and generated a fruit BRIX (Fig.3A) that resemble their nongrafted lines (Fig.2B). In treatment group, when *bip0663* served as the rootstock for Lukullus (scion/rootstock: Lukullus/*bip0663*), fruit BRIX of Lukullus was significantly increased compared with its self-grafted line (Lukullus/Lukullus), and when Lukullus served as the rootstock for *bip0663* (*bip0663*/Lukullus), fruit BRIX of *bip0663* was decreased compared with to its self-grafted line (*bip0663*/*bip0663*) (Fig.3A). In addition, measurements of fruit yield show that there was no significant difference in fruit yield between control and treatment group (Fig.3B). These results indicate changes in fruit BRIX between *bip0663* and Lukullus were either caused by photosynthate quantities or efflux from leaves or changes in root sink strength.

**Fig.3.**
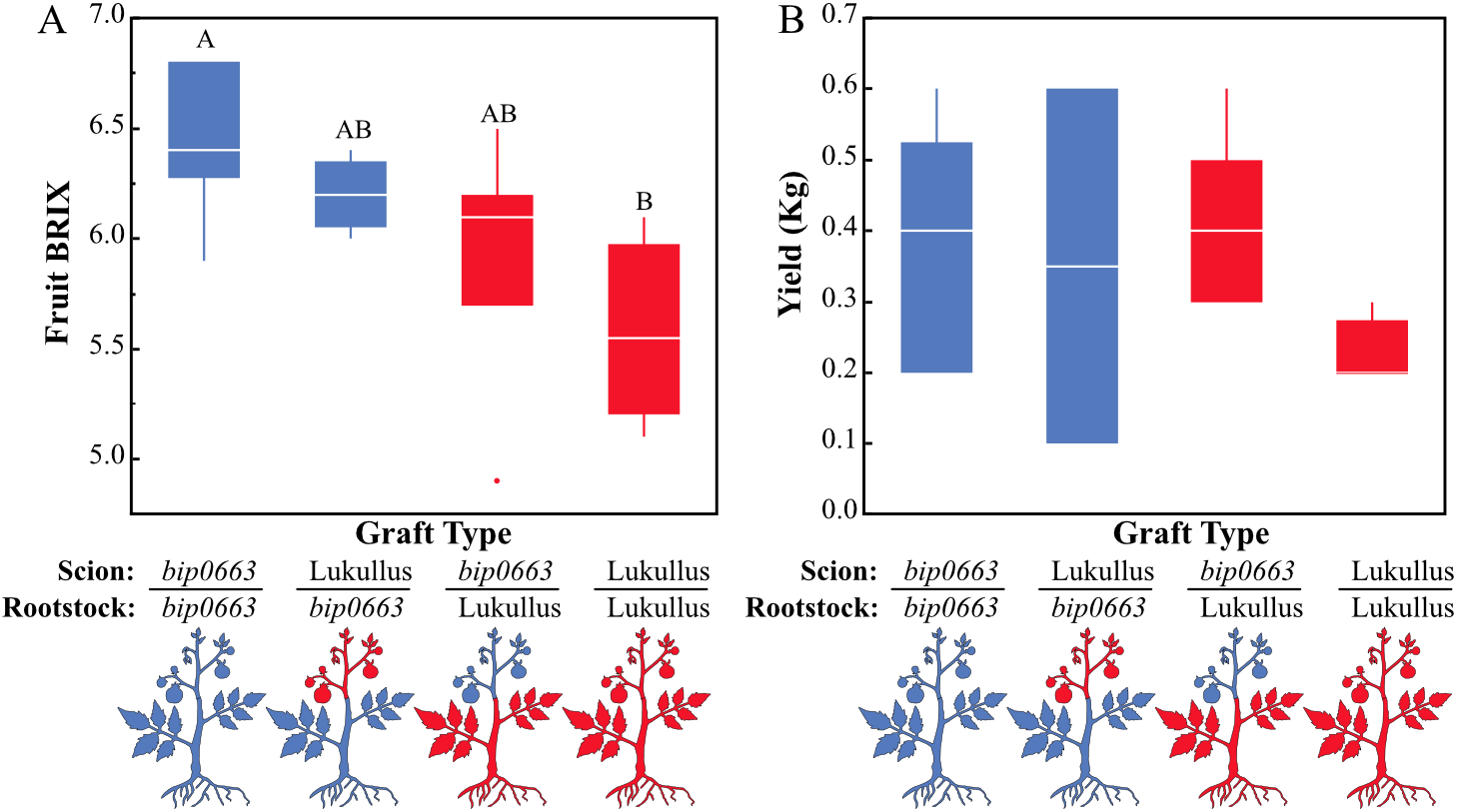
Fruit BRIX and yield of grafts. A, Fruit BRIX; B, Yield. The grafting assay used a standard least squares model of data. Graft junctions are diagramed, *bip0663* (blue) and Lukullus (red).

It was possible that the grafts containing Lukullus rootstock could have reduced fruit sugar due to increased sink strength of Lukullus roots. To determine if differences in root sink strength could have contributed to the differences in fruit BRIX, a second grafting assay was performed in which the roots of each genotype were reciprocally grafted. Root grafts with *bip0663* scion had the same high fruit BRIX and yield, regardless of the root genotype. Similarly, root grafts with Lukullus scion showed lower fruit BRIX and yield, regardless of the root genotype (Fig.4). There were no significant differences in fruit BRIX or yield when comparing *bip0663*/*bip0663* to *bip0663*/Lukullus or Lukullus/Lukullus to Lukullus/*bip0663* grafts (Fig.4). This supports the hypothesis that the presence of *bip0663* source leaves led to increased fruit BRIX of the grafts.

**Fig.4.**
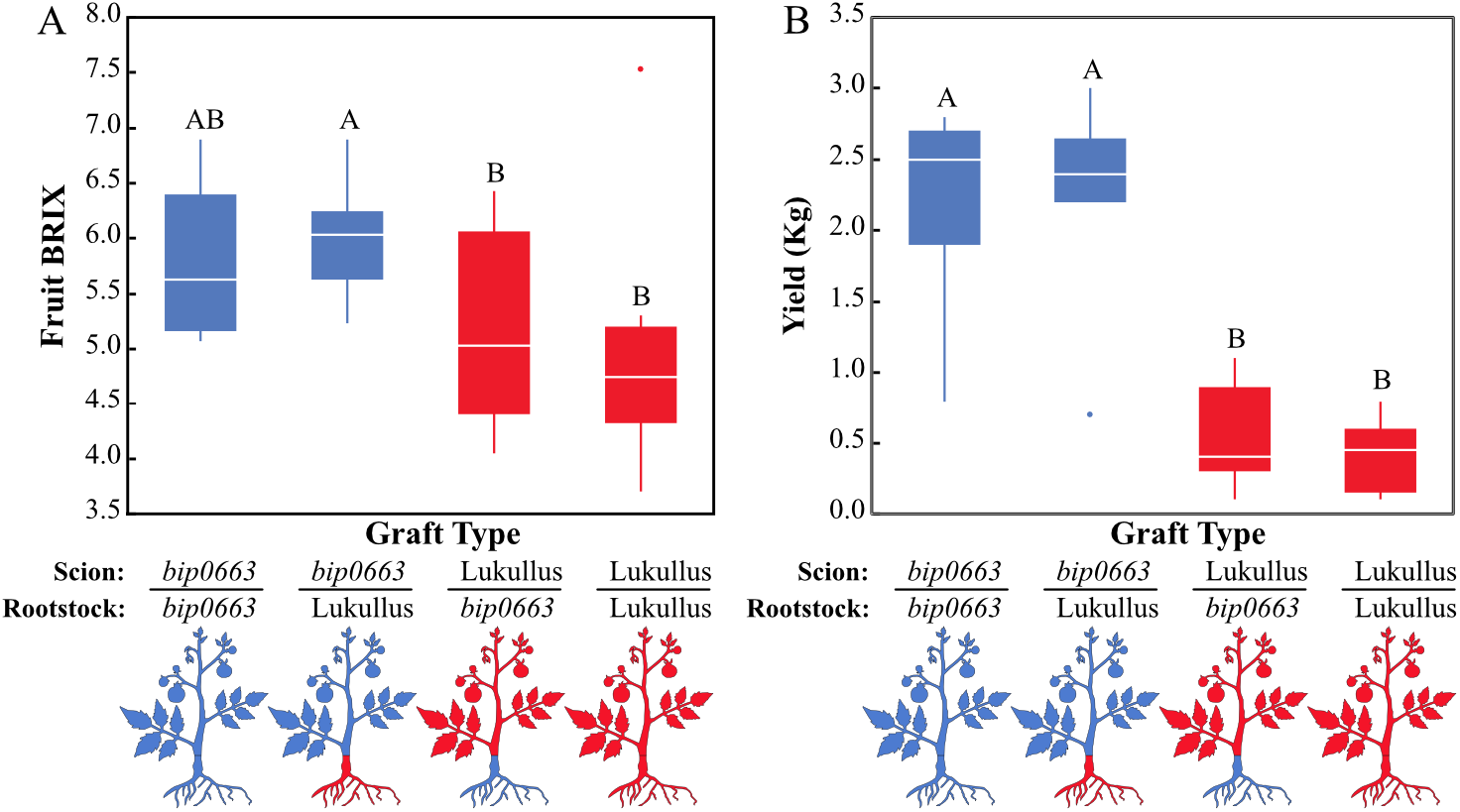
Fruit BRIX and yield of Root Grafts. A, Fruit BRIX; B, Yield. The root grafting assay used a mixed model including random effect of positional data. Graft junctions are diagramed, *bip0663* (blue) and Lukullus (red).

To further examine the differences between *bip0663* and Lukullus source leaves, the sugar and starch content of leaves was measured at dusk. The leaves of *bip0663* had less sugar (p=0.0939) and less starch (p=0.0074) than the leaves of Lukullus (Fig.5). This suggests that *bip0663* source leaves export photosynthate more efficiently than those of Lukullus.

**Fig.5.**
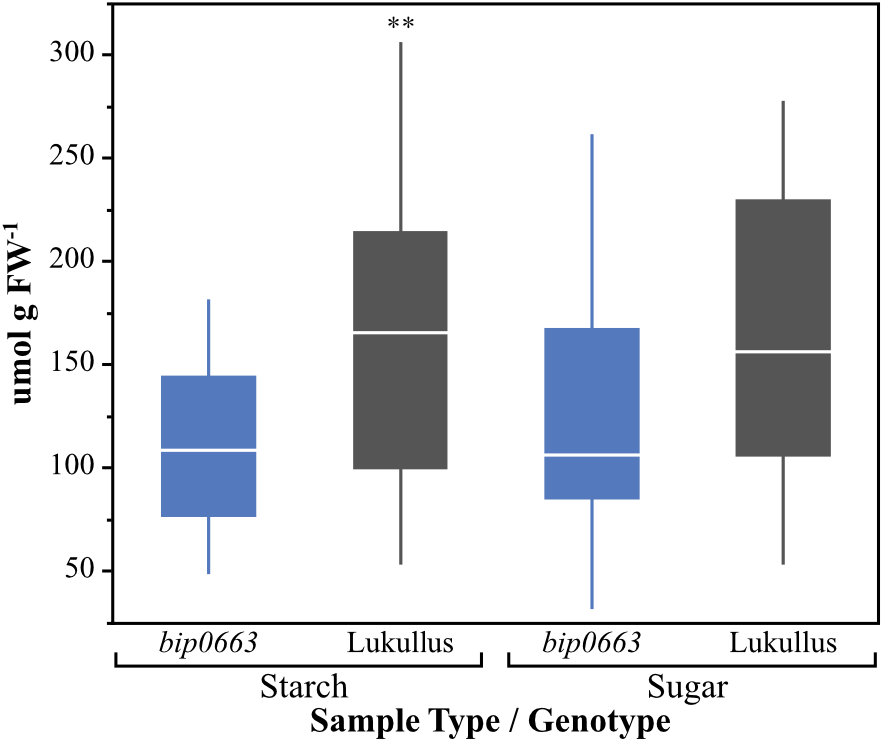
Quantified leaf sugar and starch contents for *bip0663* and Lukullus. *bip0663* has lower leaf sugar than Lukullus (p=0.0939) and significantly lower leaf starch compared to Lukullus (p=0.0074).

### Leaf shape analysis of *bip* mutants and wildtype

Since the grafting experiment indicated that changes in leaf features could drive changes in fruit BRIX, and recent studies in an introgression population and a selected group of heirloom tomato varieties show leaf shape is strongly correlated with fruit BRIX and sugar accumulation (Chitwood et al., 2013), we did leaf shape analysis on *bip* mutants and their isogenic wildtype cultivar to see whether changes in fruit BRIX between these genotypes were also correlated with variation in leaf shape. Results from PCA analysis of a total of 1500 primary leaflets showed that PC1 contributes 65% of all variation of leaflet shape, and is strongly correlated with leaflet area (or size) (R^2^=0.98), suggesting leaf size was the largest source of variation among leaflets of the 4 genotypes. PC4 contributes 3.8% of all variation and is correlation with aspect ratio with an R^2^ of 0.6 (Fig.6A and 6B). PC2, PC3 and PC5 were not correlated with any traditional shape measures, and may describe the overall variation in leaf symmetry (Chitwood et al., 2013; Rowland et al., 2019). Fig.6C show results of traditional leaf shape measurements. Both *bip2* and *bip0663* shows a similar change in leaf shape compared with their corresponding isogenic wildtype backgrounds: leaflet area and solidity decreased, and leaflet aspect ratio and circularity increased. These results indicate that mutations in the *BIP* gene not only regulate leaf complexity, but also alter leaflet shape. Lukullus and *bip0663* displayed leaflet aspect ratio and circularity significantly higher than that of *bip2* and M82. Compared to both isogenic wild genotypes, both *bip* mutants displayed rounder leaflets, with significantly higher aspect ratios and circularity measures.

**Fig.6.**
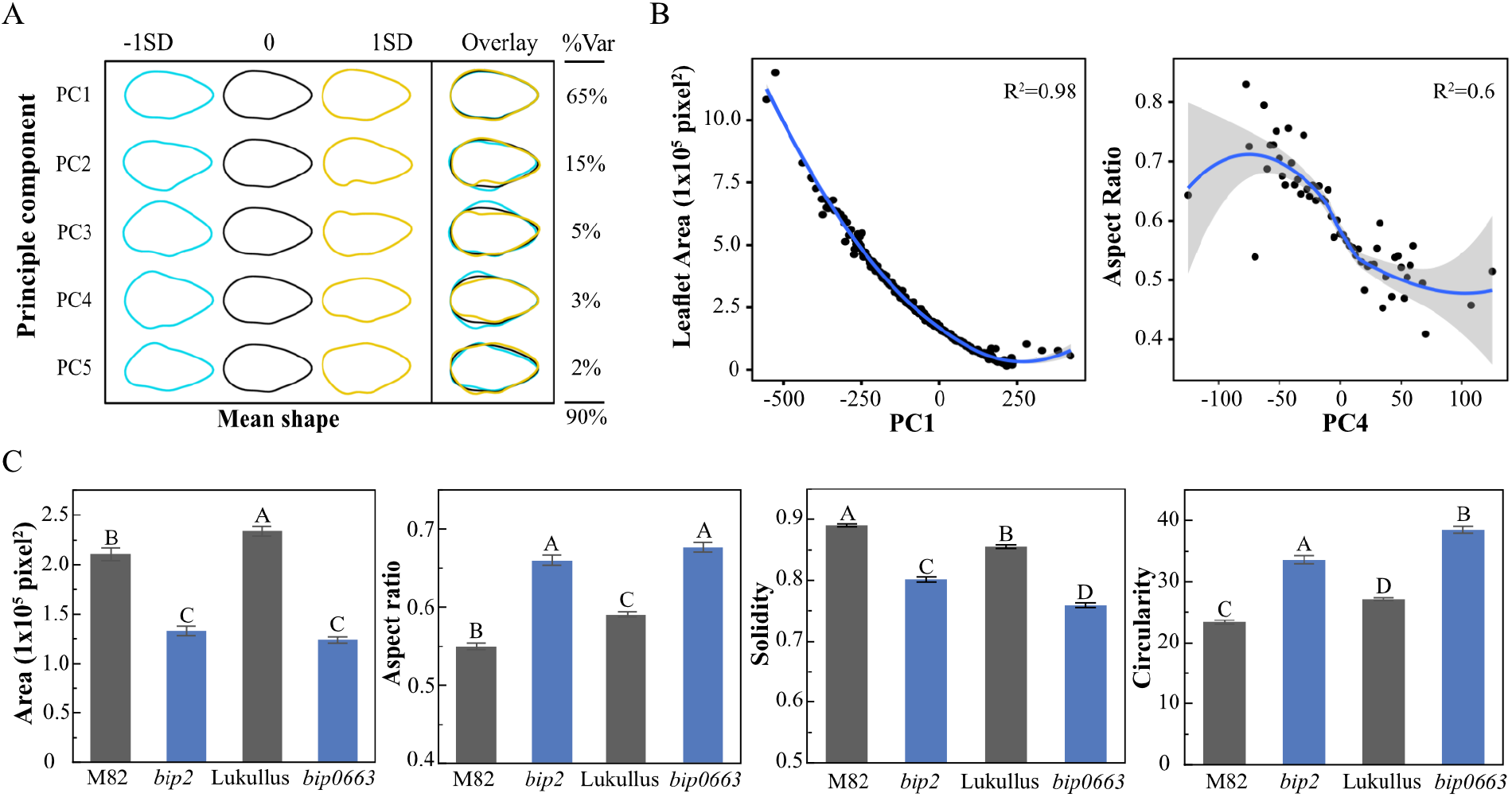
Leaf shape analysis. A and B, PCs of all primary leaflets measured (A) and their relationship to traditional shape measures (B); C, Traditional shape measures of primary leaflets of *bip2*, *bip0663*, M82 and Lukullus.

### Leaf photosynthesis of *bip* mutants and wildtype

In order to determine leaflet contribution to fruit sugar, we performed leaflet *A*, *gst*, PARi, and ΦPSII measurements on *bip* mutants and their corresponding isogenic wildtype cultivars during fruiting stage (week 17-21). Correlations between *A* and *gst* (Fig.7A), were similar between *bip2*, *bip0663*, and their corresponding wildtype cultivars, with photosynthesis reaching a maximum rate between 0.6 to 0.8 *gst.* All genotypes reached a PARi of greater than 1100 μmol m^-2^ s^-1^ over the growing season. Due to differences in determinacy between the cultivars, *bip2* and M82 reached this peak 1∼2 weeks earlier than *bip0663* and Lukullus (Fig.7B). Corresponding to increasing PARi with age, ΦPSII had an overall downward trend across the whole fruiting season in all genotypes (Fig.7C). Across the whole season, average photosynthetic rates of *bip2* and M82 were higher than those of *bip0663* and Lukullus (Fig.7D). **Leaf Vascular Patterning of *bip* Mutants and Wildtype**

**Fig.7.**
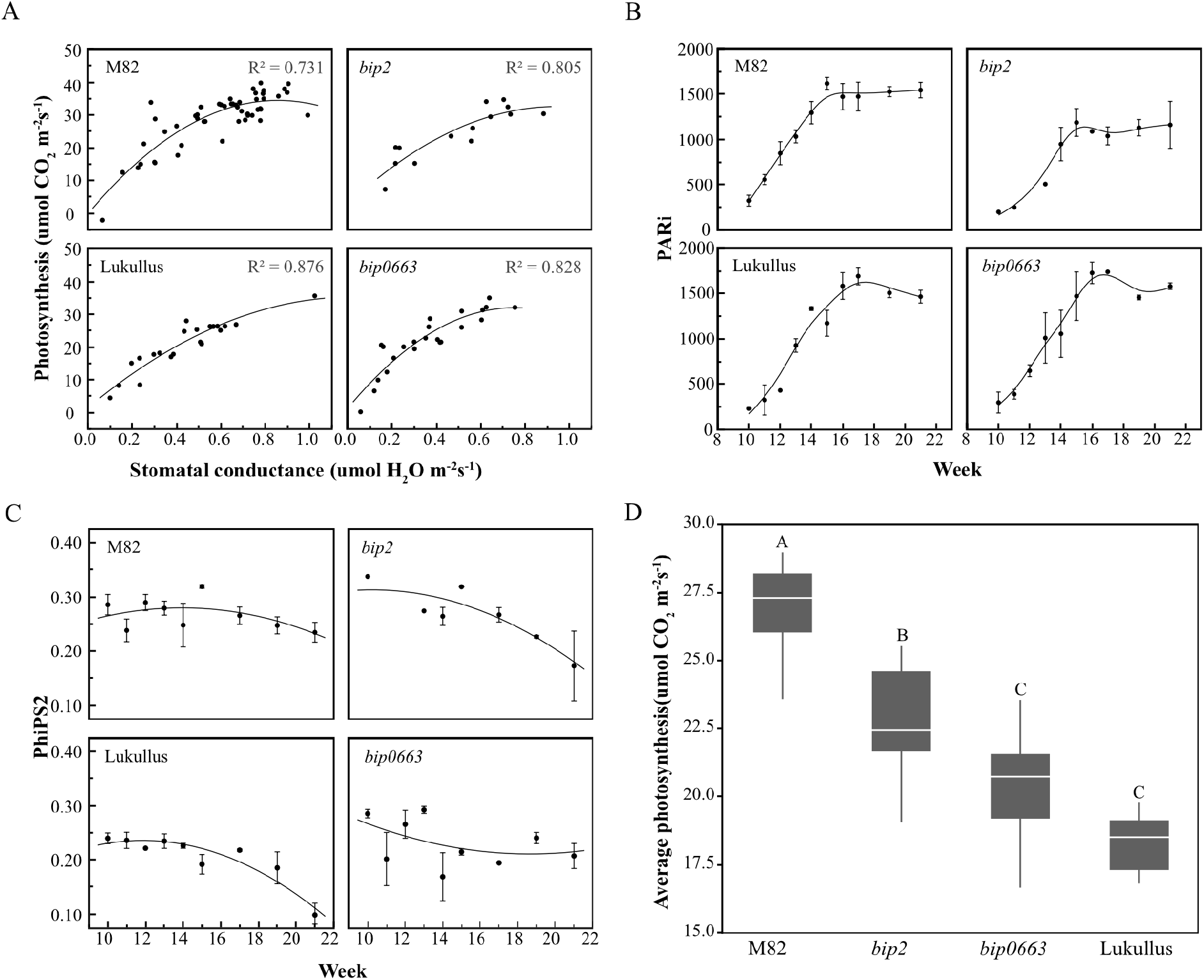
Leaf photosynthesis across the whole field season. A, Photosynthetic rate and stomatal conductance were measured weekly using the LI-6400XT across the whole field growing season. PARi (B) and phiPS2 (C) was measured weekly across the whole field growing season and values are the mean measurements over time and error bars represent standard error. D, The average photosynthetic rate across whole measured season in umol CO_2_ m^-2^s^-1^.

Generally, carbon-compounds (such as sugar) required for fruit development and ripening are primarily synthesized in the leaf and then transported to fruit by phloem (Cocaliadis et al., 2014). Therefore, we determined the leaf vascular density of each genotype (Fig.8A and Fig.8B). While the leaf vascular density of *bip0663* was significantly reduced compared with Lukullus, the leaf vascular density of *bip2* was the same as M82. In addition, overall leaf vascular density of *bip0663* and Lukullus both was lower than that of *bip2* and M82. Given that changes in leaf vascular density roughly corresponded with the difference in fruit BRIX, we performed a correlation analysis between leaf vascular density and fruit BRIX in all *bip* mutants and isogenic wildtype cultivars (Fig.8C),. A negative correlation with *R*^2^=0.844 and *p* value<2.2e-16 was detected (Fig.8C). This suggest fruit BRIX is tightly and significantly negatively correlated with leaf vascular density.

**Fig.8.**
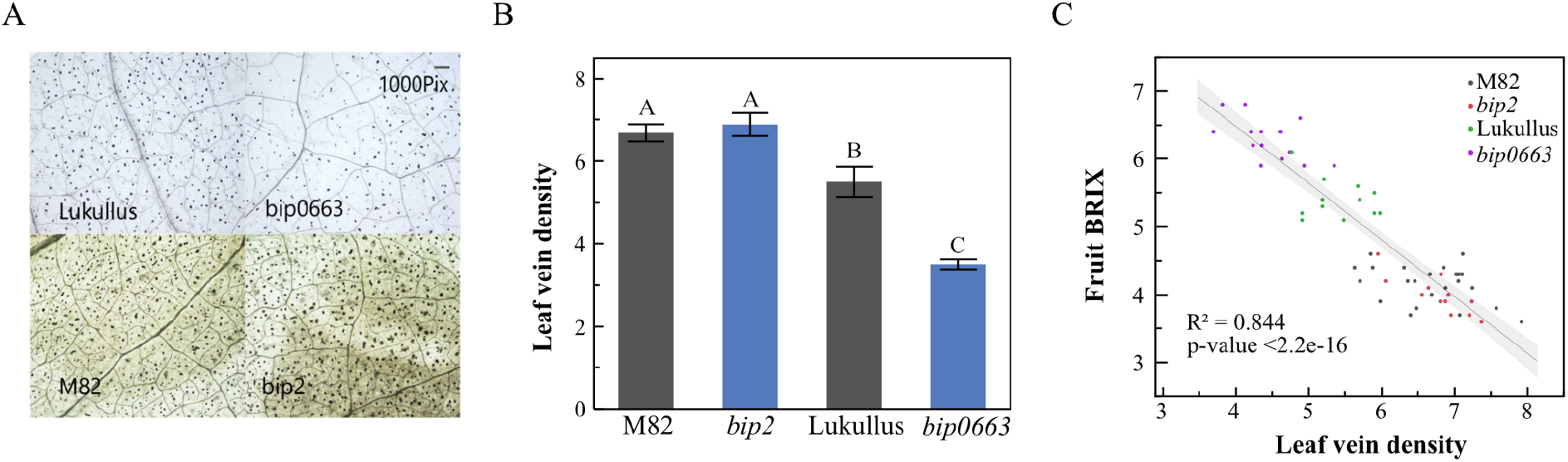
Leaf vascular patterning micrographs and density analysis A, Leaf clearing images of *bip* mutants and their isogenic wildtype cultivars, micrographs taken at a fixed magnification of 4X. B, Leaf vein density was measured by leaf GUI for each individual leaf disc and a mean value was calculated, error bars represent standard error. C, Correlation between leaf vein density and fruit BRIX across *bip* mutants and their wildtypes was analyzed using JMP with “Spearman” method.

### Fruit BRIX of *bip3* and Genotypes with Altered Leaf Vascular Density

To further confirm the correlation between leaf vascular density and fruit BRIX, we measured the leaf vascular density and fruit BRIX of several other tomato varieties (Fig.9). A mutant phenotype with high fruit BRIX, was found and identified in a mutagenized M82 line, and had rounder fruits, hence we called it M82 “Round” Morph (*MRM*). *MRM* had low leaf complexity and big leaflets similar to M82, but with fruit BRIX and yield both significantly higher than M82 (Fig.9A). Genotyping results show that the *BIP* gene sequence in this line is identical to that in M82. Leaf vascular density analysis of MRM identified extremely reduced vascular density compared to M82 (Fig.9A and **9B**). *bip3* is another *bip* mutant in the isogenic M82 background but with a similar leaf vein density as M82. *Silvery Fir Tree* (*SiFT*, an Heirloom tomato) and *sf^wl^* (*solanifolia*, *wox1* gene mutant) (Burko and Ori, 2013) both had a low leaf vein density (Fig.9A and Fig.8B). The leaf vein density of *MRM*, *SiFT* and *sf^wl^* was significantly lower than that of *bip3* and M82, while the fruit BRIX was significantly higher than *bip3* and M82 (Fig.9B **and** C). There was a high correlation (R^2^=0.855, p-value=2.2e-16) between leaf vein density and fruit BRIX, further indicating a connection between these two traits (Fig.9D).

**Fig.9.**
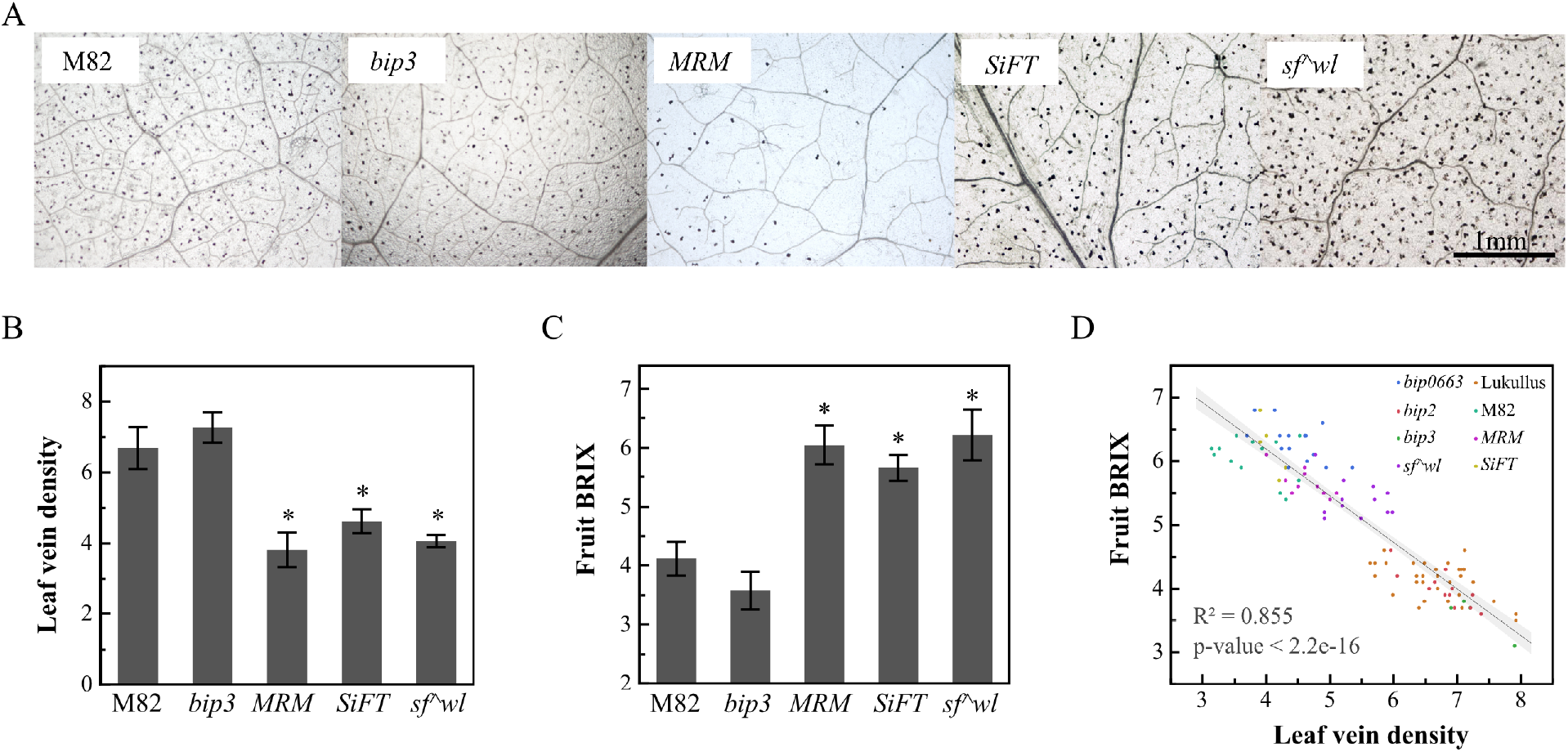
Leaf Vascular Density and Fruit BRIX in Other Tested Tomato Varieties. A, Leaf clearing images of *bip* mutant, and other varieties, taken at a fixed magnification of 4×10. B, Leaf vascular density was measured using leaf GUI for each individual leaf disc and a mean value was calculated, and error bars represent standard error. C, BRIX of tomato fruits. D, Correlation between leaf vein density and fruit BRIX analyzed using JMP with “Pearson” method.

### Differential Gene Expression

To understand global transcriptional alterations between high and low BRIX phenotypes and their possible links to leaf vascular density, we performed RNA-seq analysis on leaves of *bip0663*, Lukullus, M82, and *bip2*. RNA was extracted at three developmental stages (SAM, young leaf and mature leaf) during fruiting stage (17 weeks from germination).

Differentially expressed genes were defined using ANOVA-like test with |logFC|>1 and FDR<0.05. A total of 750 DEGs were obtained between high BRIX genotypes (HB, with low leaf vein density, included *bip0663* and Lukullus) and low BRIX genotypes (LB, with high leaf vein density, included *bip2* and M82) across three stages. Further, 205 genes were defined to be differentially expressed between *bip0663* and Lukullus, and 602 genes were defined to be differentially expressed between HB and LB. We found that 57 genes were shared between these two sets of DEGs (Fig.10A**, Table S1)**. These 57 DEGs were overrepresented in many GOslim categories including cellular process, response to stress, metabolic process, and carbohydrate metabolic process (**Table S2**).

**Fig.10.**
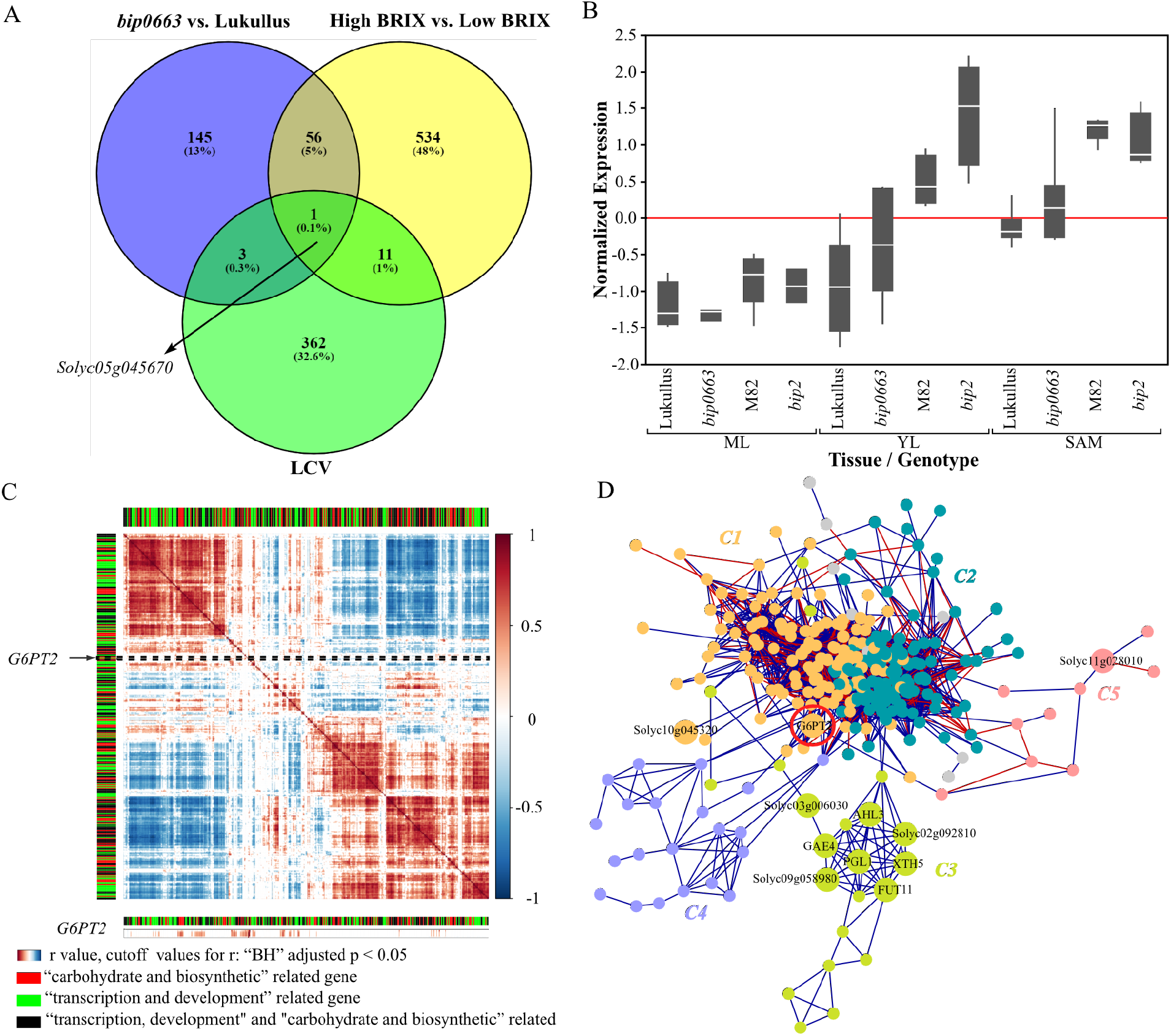
Gene Co-expression Analysis of Differentially Expressed Genes, A, Venn diagram between *bip0663* vs. Lukullus differentially expressed genes, HB vs. LB (High BRIX vs. Low BRIX group) different expressed genes, and literature-curated vascular genes (LCV). B, Normalized expression of *GPT2* (Solyc05g045670) in Lukullus, bip0663, M82, and bip2 in mature leaves (ML), young leaves (YL), and shoot apical meristem (SAM). C, Correlation analysis of DEGs enriched in “transcription and development” and “carbohydrate and biosynthetic”-related GO terms, cut off value for Pearson correlation coefficient was “BH” adjusted p value<0.05. *G6PT2* was positively correlated with many genes related to carbohydrate and biosynthetic process (| r | value>0.5 and “BH” adjusted p value <0.05). D, Gene co-expression network constructed across *bip0663*, Lukullus, *bip2* and M82 using DEGs enriched in “transcription and development” and “carbohydrate and biosynthetic processes”-related GO terms. Nodes (genes) with at least one edge (298 nodes and 7517 total edges) are represented. The communities are represented by different colored nodes (*C1*-*C5*). Top DiffCorr genes (Table S5) are labeled in black, *G6PT2* is circled in red.

Many genes regulating developmental related processes exhibit broadly conserved function across plant species, and this is particularly true of genes relating to leaf veins and leaf development (Townsley and Sinha, 2012). Therefore, to narrow down target genes that may be causal for leaf vascular density alterations, we focused on literature curated genes that are already known to regulate leaf vein development in the model species *Arabidopsis thaliana* (Wenzel et al., 2008; Paul et al., 2013, Parsons-Wingerter et al., 2014). Tomato genes that are orthologous to literature curated genes in *Arabidopsis thaliana* were generated and named LCV (the literature-curated vascular related genes, **Table S3**). Interestingly, a shared LCV gene *Solyc05g045670* (*GLUCOSE-6-PHOSPHATE/PHOSPHATE TRANSLOCATOR 2*, *GPT2*) was detected in both sets of DEGs (Fig.10A). In SAM and YL, expression of *GPT2* were down regulated in both *bip0663* and Lukullus compared to other genotypes, and the expression of *GPT2* in *bip0663* was down regulated relative to Lukullus (Fig.10B). Another genotype identified to have low vascular density and high fruit BRIX, *SiFT* (Fig. 9A **and** B), also has down regulated *GPT2* expression compared to M82 in P4 leaf primordia (**Figure S3**) (Nakayama et al., 2020). *Solanum pennellii*, a wild relative of tomato, with high fruit BRIX, has lower expression of *GPT2* in leaves compared to M82 (Koenig et al., 2013). Analysis of its promoter sequence compared to M82 shows SNPs in several putative cis-regulatory elements identified by NSITE-PL (Shahmuradov et al., 2015), including the transcription factor binding sites for HVH21 and GT1 (LRE), and the G-box binding site (**Figure S4**) (Krusell et al., 1997; Hiratsuka et al., 1997; Ezer et al., 2017).

### Co-expression analysis of differential expressed genes

To detect changes in the patterns of expression of the DEGs between phenotypes with different fruit BRIX, we performed DiffCorr analysis (Fukushima, 2013; Ichihashi et al., 2014) on all detected DEGs. DiffCorr networks were separately constructed on all DEGs between *bip0633* and Lukullus comparison (bipvsLu Diffcorr network, **Figure S5A**) and also between HB and LB comparisons (HvsL Diffcorr network, **Figure S5B**) with cutoff adjusted p-value<0.05. In these DiffCorr networks nodes with more connected edges represent genes which have the most differential co-expressions between the two compared co-expression networks. Of the top 20 ranked Diffcorr genes, 11 genes were shared and highly differentially co-expressed between bipvsLu and HvsL Diffcorr networks (**Figure S5 and Table S4).** The expression patterns of these genes were highly different between *bip0663* and Lukullus, and also between HB and LB comparisons. Six of these 11 genes were reported to be involved in regulating cellular processes, cell walls, leaf vein development or sugar metabolism. During leaf development, *LTP4* (*Non-specific lipid-transfer protein*, *Solyc08g067500*, leaf vascular development related), *AHL3* (*AT hook motif DNA-binding family protein*, *Solyc03g007150*, leaf vascular development related) and *FUT11* (*Alpha 1,3 fucosyltransferase*, *Solyc01g108110*, cell, cell wall, and leaf vascular development related) were all down regulated in *bip0663* and Lukullus, in young leaves and mature leaves compared with *bip2* and M82 (Wenzel et al., 2008; Zhou et al., 2013; Bakker et al., 2001). *PGL1* (*6-phosphogluconolactonase*, *Solyc01g010250*, cell, cell wall, and leaf vascular development related) was up regulated in *bip0663* and Lukullus in ML (**Figure S6)**. Thus, these genes might play an important role in regulating fruit BRIX and leaf vascular density between *bip0663*, Lukullus, *bip2*, and M82.

In addition, many DEGs detected between the HB and LB groups were involved in regulating both development and carbohydrate metabolism (**Table S5**). We therefore hypothesize that these genes might be co-expressed during leaf vascular development and involved in regulating leaf sugar metabolism either directly or indirectly through vascular transport processes. To validate this hypothesis, we performed correlation analysis and constructed a gene coexpression network across all genotypes using DEGs enriched in “transcription and development” and “carbohydrate and biosynthesis” related GO terms. Correlation analysis showed most of “transcription and development” related DEGs and “carbohydrate and biosynthesis”-related genes are highly correlated, whereas *GPT2* was shown to be positively correlated with many genes related to carbohydrate and biosynthetic process (**Fig.10C** and **Table S6**, Pearson correlation with BH adjusted p value<0.05), such as *Solyc01g074030* (Beta-glucosidase gene). Interestingly, *LTP4* showed significantly positive correlation with *GPT2*. In the Co-expression network, community *C3* shows significant enrichment of the shared top ranked DiffCorr genes (including *AHL3*, *PGL1*, *XTH5*, *GAE4* and *FUT11*) listed in Table S5 and is connected to other two communities (*C1* and *C2*), which compose a core network (with hub genes with >100 edges). *C1* was enriched for genes related to transcription initiation and cell fate specification GO terms, and *C2* contained genes involved in glucose metabolic process, carbohydrate biosynthetic process, plant organ development, hormone-mediated signaling pathway and shoot system development (**Table S7)**. *GPT2* can be found on the border of the core network, within *C1* and co-expressed with several genes with hubs >50, revealing it might be an important gene that connects leaf vein development and carbohydrate biosynthetic processes together. *C3* contained genes involved in glucose-6-phosphate metabolic process, and *C5* was enriched for genes involved in cellular carbohydrate biosynthetic processes.

## DISCUSSION

Since sugars synthesized in leaves are the primary source of energy for fruit development and ripening (Cocaliadis et al., 2014), we and others have investigated the link between leaf and fruit. Previous studies have shown leaf traits, such as leaf shape and size are strongly correlated with fruit quality and yield (Chitwood et al., 2013; Rowland et al., 2019, Gupta et al. 2020). In this study, we used grafting experiments to analyze the influence of leaf traits on fruit sugar level. Our results show that changes in leaves can cause variation in fruit BRIX. Leaf shape analysis showed that both *bip* mutants had rounder leaflets, with significantly higher aspect ratios and circularity measures, compared to both isogenic cultivars. This suggests *bip* mutations caused not only high leaf complexity but also rounder leaves. Results also show that leaflet roundness of *bip0663* and Lukullus (HB genotypes) was significantly higher than *bip2* and M82 (LB genotypes). These results agree with previous research that showed tomato with rounder and more circular leaves tend to have the highest sugar content in their fruit (Chitwood et al., 2013; Rowland et al., 2019). Although *bip2* has rounder leaves than M82 it does not have measurably higher fruit BRIX (Fig. 2B), suggesting that the roundness caused by the *bip* mutation does not, by itself, lead to increased BRIX.

A previous study (Chitwood et al., 2013) suggested the correlation between leaf shapes and fruit sugar content may be due to the impact of leaf shape on photosynthetic capacity. However, we found that across the whole season, average photosynthetic rates of *bip2* and M82 (LB genotypes) were higher than those of *bip0663* and Lukullus, perhaps resulting from high vein density (vein length per unit area) in the LB genotypes (Sack and Scoffoni, 2013). Thus, the variation in fruit BRIX between these genotypes was likely due to aspects other than leaf photosynthesis. In addition, while mutations in the *BIP* gene cause increased leaf complexity, only *bip0663* has increased fruit BRIX compared with Lukullus. This suggests that leaf complexity is not the cause of changes in fruit BRIX between *bip0663* and Lukullus. This is consistent with the results from path modeling connections between leaf traits and fruit BRIX in multiple heirloom cultivars (Rowland *et al.*, 2019). The same path modeling also indicated that photosynthesis was not a major contributor to fruit BRIX (Rowland *et. al.*, 2019).

In leaves, sugars synthesized through photosynthesis are first loaded into leaf veins and then transported out of the leaves to the rest of the plant sinks, such as fruit. Thus, leaf veins also play an important role in fruit sugar accumulation by contributing to transport of carbohydrates (Cataldo, 1974; Haritatos et al., 2000; Sack and Scoffoni, 2013). Theoretically, higher leaf vein density will increase the contact between vascular and photosynthetic tissues and lower the distance of photosynthate transportation (Adams et al., 2007; Sack and Scoffoni, 2013). Several studies have shown high leaf vein density can not only enable higher *K*leaf (leaf hydraulic conductance) and higher rates of gas exchange per leaf area (Sack and Frole, 2006; Boyce et al., 2009; Brodribb et al., 2010), but also can improve phloem loading (Russin and Evert, 1984). Based on these studies, with all else being equal, plants with high leaf vein density should have higher fruit sugar levels. However, in this study leaf vein density was found to be negatively correlated with fruit BRIX in *bip* mutants and their corresponding isogenic cultivars. This may be caused by the following factors: Leaves with low leaf vein density can enhance mesophyll light capture in shade and result in increased photosynthate (Brodribb *et al*. 2007; Zhu *et al*., 2012; Sack and Scoffoni, 2013). Additionally, many free-ending veins were observed in *bip0663* and Lukullus leaves (Fig.9), which can promote sugar export (Adams *et al*., 2007).

Leaf veins, similar to leaf shapes, also have a remarkable diversity in architecture within and across species. Numerous studies on mutant phenotypes have shown auxin signaling networks play an important role in patterning leaf vasculature. Polar auxin transport (PAT), as a positional cue, determines the sites of vascular cell differentiation, and inhibition of auxin transport can lead to excessive leaf vein growth in Arabidopsis (Enrico *et al*., 2006). Since auxin also patterns blade outgrowth, leaf morphogenesis and vascular pattern formation can be coupled (Koenig et al., 2009). Genetic and molecular studies have identified many important genes that are involved in regulating vascular pattern formation that are related to auxin transport or biosynthesis (Miyashima et al., 2013; Biedroń and Banasiak, 2018). For example, *MONOPTEROS/AUXIN RESPONSE FACTOR 5 (MP*/*ARF5)* and *TARGET OF MONOPTEROS 5* (*TMO5)* are essential for provascular establishment (Han *et al*., 2018), *AT HOMEOBOX 8* (*ATHB8*) defines preprocambial cell state that is a prelude to vein formation, and *ACAULIS5* (*ACL5*) and *BUSHY AND DWARF 2* (*BUD2)*, which regulate polyamine synthesis, can inhibit auxin-induced xylem differentiation (Baima et al., 2014), while *PHLOEM INTERCALATED WITH XYLEM* (*PXY*) and *CLAVATA 3/ESR-RELATED 41/44* (*CLE41*/*44)* regulate vascular organization (Etchells and Turner, 2010). Moreover, few genes such as *VASCULAR-RELATED NAC-DOMAIN 6/7* (*VND6*/*7)*, *NAC SECONDARY WALL THICKENING PROMOTING FACTOR 1/2* (*NST1*/*2)* and *ALTERED PHLOEM DEVELOPMENT* (*APL)* are also reported to be related to leaf vascular patterning though regulation of phloem or xylem differentiation (Bonke et al., 2003; Mitsuda et al., 2007; Yamaguchi et al., 2010). In addition, using auxin transport inhibitor (NPA) induced vascular overgrowth, Wenzel and coworkers (Wenzel et al., 2008) identified many vascular related genes that were previously not known to have a role in vascular differentiation in Arabidopsis. These include *At1g61800* (*GLUCOSE-6-PHOSPHATE/PHOSPHATE TRANSLOCATOR 2, GPT2*), *At3g16360*, *At1g07430* (*PROTEIN PHOSPHATASE 2C*), and *At5g17220* (*TRANSPARENT TESTA 19*, *TT19*). These genes had vascular-related expression in transgenic *Arabidopsis* plants and were notably up-regulated in tissue with vascular overgrowth. In our study, *GPT2* (*Solyc05g04567*, a homolog of *At1g61800*) was significantly down regulated in low leaf vein density genotypes such as *bip0663* and Lukullus, suggesting correlation between leaf vein density and expression of *GPT2*. Recent studies have shown that GPT2 functions in promoting cell proliferation, which would be one way to link GPT2 with leaf development and vascular patterning. Mutants in *gpt2* may have accelerated chloroplast differentiation (Van Dingenen *et al*., 2016). GPT2 is also involved in starch biosynthesis (Bakker et al., 2001; Dyson et al., 2015). In leaves, *GPT2* is responsible for transporting of Glucose-6-phosphate (G6P) into plastids where G6P can be used as a carbon source for starch biosynthesis. Therefore, down regulation of *GPT2* might decrease transport of glucose into plastids and increase transport of glucose out of the leaf in *bip0663* and Lukullus compared with *bip2* and M82. Our study showed decreased leaf starch and sugar in *bip0063* when compared to Lukullus, which may also be indicative of increased export of sugars from the leaf.

There are two main explanations for the relationship between leaf morphology and fruit sugar content. The first is the purely developmental link between the leaves and the fruit. Studies have shown similarity in morphology (such as size, aspect ratio, etc.) between leaves and fruits (Xiao *et al*., 2008; Chitwood *et al*., 2013). This is borne out by the fact that the morphological development of fruits and leaves is regulated by a similar gene regulatory network (Barkoulas et al., 2008). The second is that either genetic changes in overall metabolism alter leaf development or changes in carbon metabolism induce morphological changes in leaves (Hackel et al., 2006; Lawson et al., 2006; Raines and Paul, 2006). For example, *ARF4* not only regulates the morphology of leaves and fruits (Yifhar et al., 2012), the accumulation of chloroplasts and the greening of fruits (Jones et al., 2002), but was also recently shown to have influence on fruit sugar levels (Sagar et al., 2013). Because GPT2 has been shown to regulate cell proliferation and is also involved in starch biosynthesis, it could be a common regulator for leaf vein density and carbon allocation. In addition, our analyses show that DEGs enriched in “transcription and development” – GO terms were highly correlated with those enriched in “carbohydrate and biosynthetic”-GO terms. In our gene co-expression network, *GPT2* was in the core network involved in biosynthetic and development process and co-expressed with several highly connected genes (hubs >50), suggesting that it might be an important and conserved gene that connects leaf vein development and carbohydrate biosynthetic process.

## METHODS

### Plant Materials

The classic *bip* mutants used in this study and are *bip2* (isogenic cultivar: M82) and *bip0663* (*bip* accession number LA0663) (isogenic cultivar: Lukullus). M82, *bip0663*, and Lukullus seeds were obtained from the Tomato Genetics Resource Center (TGRC, http://tgrc.ucdavis.edu/). *bip2* (accession number e0652b) was obtained from the saturated mutation library of tomato (Menda et al., 2004).

### Seed germination and plant growth Conditions

Tomato seeds were treated with 50% bleach for 10 min and rinsed 3-5 times with water, then placed on water dampened Phytatrays (Sigma Aldrich). Seeds were moved to the dark and incubated at room temperature for 3 days, then transferred to a growth chamber set at 25°C with 16:8 photoperiod until seedlings had expanded cotyledons (approximately 4-7 days). The seedlings were then transplanted to 72 Seedling Propagation trays and grown in the chamber for 7 days. After that, seedlings were transferred to the greenhouse or grown for 2 weeks and then transplanted to field. The greenhouse plants were watered from the top to encourage hardening. Field plants were watered with furrow irrigation once weekly.

### Analysis of leaf complexity and shape

Mature fully expanded leaves from adult nodes (leaf 5 and above) were used for leaf complexity and shape analysis, and at least five leaves were collected from each plant for analysis. Leaf complexity is defined as the number of all leaflets present on the leaf. For leaf shape analysis, intercalary and secondary/tertiary leaflets were ignored due to their irregular shapes. Leaf shape was analyzed using a method previously described (Ichihashi et al., 2014). After leaf complexity was measured, the leaflet images were used for shape and size analysis. Leaflets were imaged using Epson Perfection V600 Photo scanner (Epson America Inc., CA, USA), and each individual leaflet image was saved as a binary image so that the leaflet was black on a white background. The binary images were then processed in R using MOMOCS (Bonhomme et al., 2014). After importing and aligning along their axes, leaflet images were then processed using elliptical Fourier (eFourier) analysis based on the number of harmonics calculated from the MOMOCS package (Bonhomme et al., 2014; Rowland et al., 2019). Traditional leaflet shape measures such as leaflet area (size), solidity, circularity, and roundness (aspect ratio) were measured based on figure pixel. PCA analysis was performed on eFourier results and statistical correlations between Principal Components (PCs) and traditional leaflet shape measures were used to determine the leaf characteristics captured by each PC.

### Determination of fruit sugar content

To measure the fruit BRIX, approximately nine tomato fruit from each plant were collected and taken to the lab where the juice was collected and BRIX measured using a refractometer (HI 96801 Refractometer, Hanna Instruments Woonsocket, RI).

### Analysis of leaf vein density

For leaf vein analysis, leaf discs with an area of 0.28 cm^2^ and were collected by a hole puncher from the second lateral primary leaflets of each plant (the sampling sites were located between second-order veins, seen in **Figure S1A**). Leaf discs were cleared using a modified method from the Ainsworth lab (Bishop et al., 2018; Rowland et al., 2019). Leaf discs were heated in 80% EtOH for 20 minutes at 80℃ and this process repeated twice or until leaf discs turned white. Leaf discs were then placed in 5% NaOH and heated to 80℃ for 5 minutes and were cooled by incubating at room temperature for 10 minutes. After that NaOH was removed and leaf discs were treated with 50% bleach for approximately 30 seconds. Bleach treatment was repeated until leaf discs were clear white. Then leaf discs were washed by ddH2O and vacuum infiltrated with 50% glycerol for 20 minutes. Cleared leaf discs were placed on slides and leaf veins of leaf disc were imaged using Eclipse C1 plus microscope (Nikon Instruments Inc., NY, USA) at a fixed magnification (4X). Leaf veins of images were traced and measured by LEAF GUI (Price, 2012), a tool that facilitates improved empirical understanding of leaf veins structure (http://www.leafgui.org). Leaf vein density is measured as leaf vein length per observed leaf area.

### Plant photosynthesis

Leaflet *A* (photosynthesis), *gst* (stomatal conductance), transpiration, and ΦPSII (percentage of photons entering PSII) were measured weekly using the LI-6400 XT Portable Photosynthesis System (LI-COR, NE, USA) from week 10 through week 15, week 17, and week 18-21(fruiting stage). Before measurements were taken, leaflets used for measuring were equilibrated for 2-3 minutes to stabilize photosynthetic rates (Rowland et al., 2019). The measurements were done on terminal leaflets. Light within the chamber was at 2000 µmol m^-2^ s^-1^ PAR, CO2 concentration within the chamber was set at 400 µmols mol^-1^, and air flow volume within chamber was 400 µmols s^-1^. Humidity, leaf, and chamber temperature (not allowed to exceed 36°C) were allowed to adjust to ambient conditions. The amount of intercepted Photosynthetically Active Radiation (PARi) was also measured in four orientations per plant and then the average PARi was calculated (Rowland et al., 2019).

### Grafting assay

The grafting assay used self-grafting as controls and reciprocal grafting of the two genotypes. Lukullus (scion) grafted on *bip0663* (rootstock), and *bip0663* (scion) grafted on Lukullus (stock) at the first internode as the treatments, while Lukullus grafted on Lukullus and *bip0663* grafted on *bip0663* at the first internode served as controls (experiment setting seen in **Figure S1B-D**). Thirty-day old plants grown in the chamber were used for grafting. The grafted plants were allowed to recover for about 7 days before being transferred outside into 2-gallon pots. Once the grafts start flowering, leaves from scion and flowers from the stock were continuously removed to ensure that growth of fruit on scion was only driven by root stock leaves as source. The vegetative biomass, leaf shape, leaf complexity and leaf veins (rootstock), fruit yield, and fruit sugar (scion) were measured.

### Root Grafts

For the root grafting control group, *bip0663* scions were grafted onto *bip0663* rootstocks, and Lukullus scions were grafted onto Lukullus rootstocks. For the treatment groups, *bip0663* scions were grafted onto Lukullus rootstocks and Lukullus scions were grafted onto *bip0663* rootstocks. This grafting was done with 30 day-old plants, and the junction for these grafts was in the internode between the cotyledons and first true leaves. These grafted plants were allowed to recover for 2 weeks before being transferred outside into 2-gallon pots. The grafts were allowed to flower and fruit freely and were regularly checked to assure that there were no leaves emerging from the rootstock portion of the grafts. This allowed us to determine if differences in root sink strength could lead to differences in growth of fruit. The yield and fruit sugar were measured.

### Leaf Sugar and Starch Measurements

In order to determine the sugar and starch content of *bip0663* and Lukullus, leaf discs with an area of 0.28 cm^2^ and were collected with a hole puncher from the second lateral primary leaflets of each plant (the sampling sites were located between second-order veins, seen in **Figure S1A**). These samples were taken at dusk and immediately placed into microcentrifuge tubes containing 500ul of 100% EtOH. Samples were heated at 80°C for 20 minutes. Supernatant was immediately removed, stored at −20°C, and used for determination of sugar content (including sucrose, glucose, and fructose). Sugar content was determined as previously described (Rowland et al., 2019). Leaf discs were kept for further processing and starch extraction. Residual sugar was removed from leaves by two additional rounds of heating in 500ul 100% EtOH, discarding the supernatant each time. Leaf discs were resuspended in 500ul 5% NaOH and heated to 80°C for 20 minutes. Samples were cooled and neutralized with 125ul 5M HCl. The supernatant was removed, and leaf discs were rinsed 2X with ddH2O. Discs were resuspended in 500μL 50mM Sodium Acetate Buffer and bead beaten by hand for 30 seconds after the addition of a small metal bead. Samples were centrifuged for 1 minute at 13K rpm. 25ul of a starch degradation mixture, containing a final concentration of 1.3U Amyloglucosidase and 220U α-Amylase, was added. Samples were incubated at RT for 1 hour, followed by 65°C for 23 hours. Enzymes were deactivated by heating to 80°C for 5 minutes. These samples, now containing glucose were quantified using the same protocol as for sugar samples above.

### RNA Extraction

The shoots or leaf primordia were sampled and frozen in liquid nitrogen or stored at −80°C. About 100 mg tissue was ground using a Bead-Beater (Bio Spec Products Inc., OK, USA) and RNA was then extracted using a published protocol in routine use in the laboratory (Townsley et al., 2015).

### RNA-seq Library Preparation and Sequencing

Tissue (e.g. shoot apical meristem, young leaf, and mature leaf) used for RNA-seq libraries for Illumina sequencing were collected using the method previously described (Townsley et al., 2015). RNA-seq libraries were prepared from collected tissues using the BrAD-seq method (Townsley et al., 2015). Libraries were prepared from 5 replicates of each type of tissue, collected at fruiting stage (week 17). These RNA-seq libraries were sequenced at the Vincent Coates Genomics facility at University of California, Berkeley on a single lane of the Illumina Hi-Seq 4000 platform and 50-bp single-end reads were generated. A total of 326 M raw paired-end 100 bp reads were generated, ranging from 5.7 to 16.3 M reads per library.

### Preprocessing of Illumina Reads

Illumina Reads were trimmed and mapped to reference using CLC Genomics Workbench 11 (https://www.qiagenbioinformatics.com/). Low-quality reads with average Phred quality score <20 and low-quality bases from 3’ end of the reads were trimmed using the Trim tool in CLC Genomics Workbench. The clean reads were aligned to the tomato reference genomic sequence (ITAG 3.0, *Solanum lycopersicum* Heinz 1706) by using the Large Gap Read Mapping tool in CLC Genomics Workbench, which models the presence of introns in the tomato genome reference sequence, but are expected to be absent from the corresponding transcriptome. We used the Transcript Discovery Plugin in CLC Genomics Workbench to generate a consensus CDS mapping track based on the existing reference genome (ITAG3.2; https://solgenomics.net). The new transcripts tracks were used as reference for subsequent read mapping.

### Differential Expression Analysis

Reads from individual libraries were mapped to the annotated transcripts reference using default parameters in the RNA-seq mapping tool in CLC Genomics Workbench. Then differential expression analysis between *bip* mutants and wildtype across three tissues was carried out using limma-voom (Law et al., 2014; Costa-Silva et al., 2017). Transcripts with very low estimated counts (cpm<2) were removed using limma-voom. Genes with |logFC|>1 and FDR<0.05 were considered to be differentially expressed genes (DEGs).

### Coexpression and DiffCorr network analysis

DEGs detected between *bip0663* and Lukullus, and between low (M82 and *bip2*) and high (Lukullus and *bip0663*) BRIX phenotypes were used for constructing gene co-expression networks with the cutoff values for Pearson correlation coefficient (adjusted P < 1.0×10^−8^) to capture gene interactions (Chitwood et al., 2013). Networks were constructed using the WGCNA package in R, and network communities were determined by the Fast-Greedy modularity optimization algorithm. Network visualization was done in Cytoscape (Shannon et al., 2003). Gene ontology of network communities was explored using PANTHER GO-Slim (Mi et al., 2019).

DiffCorr analysis was performed on all detected DEGs using the DiffCorr Bioconductor package in R (Fukushima, 2013; Ichihashi et al., 2014). In addition, a gene co-expression network across *bip0663*, Lukullus, *bip2* and M82 was constructed using DEGs enriched in “transcription and development” and “carbohydrate and biosynthetic”-related GO terms.

### GO Enrichment Analysis

GO enrichment analysis of the differentially expressed genes was performed using the GOSeq Bioconductor package in R (Young et al., 2012). GO analysis for gene co-expression network communities was performed using PANTHER (Thomas *et al*., 2003; Mi et al., 2019).

### DNA extraction and DNA-seq library preparation

DNA was extracted from 4 week old M82 plants using GeneJET Plant Genomic DNA Purification Mini Kit (Thermo Scientific, Waltham, MA, USA). DNA-Seq libraries were prepared based on BrAD-seq (Townsley et al., 2015) with the following modifications: After DNA fragmentation with Covaris E220 (Covaris, Inc. Woburn, MA, USA), the fragmented DNA was end-repaired, A-tailed, and adapter ligated with Y-adapter. Enrichment PCR was then performed with the adapter ligated product as described Townsley et al., 2015. After final library cleanup with AMPure beads (Beckman Coulter, Brea, CA, USA), DNA-Seq libraries were sequenced at Novogene (Novogene Inc. Sacramento, CA, USA)

### WGS Mapping and promoter analysis

*Solanum pennellii* LA01716 paired-end Illumina reads were acquired from the European Nucleotide Archive (http://www.ebi.ac.uk/ena/; accession number PRJEB5235). *S. pennellii* and M82 reads were mapped to ITAG 3.0, *Solanum lycopersicum* Heinz 1706 genome using BWA (Li et al., 2009). Post-processing of alignment was completed with Samtools (https://github.com/samtools/samtools), and VCF files were generated using Bcftools (http://samtools.github.io/bcftools/bcftools.html). Mapping and VCF files were imported into CLC Genomics Workbench 11.0 software for comparison with M82 sequence. A 3kb region upstream of *GPT2* (Solyc05g045670) was analyzed for promoter motifs using NSITE-PL (Shahmuradov et al., 2015).

### Statistical Analysis

Statistical analyses of leaf shape, leaf complexity, and leaf vein density were all performed using JMP (JMP Pro 14.0.0, 2018 SAS Institute Inc.) software. General Linear Regression Model (GLM) and One-Way ANOVA followed by Tukey’s-HSD were used to determine statistical significance in measurements. The BRIX and yield measurements from grafts were analyzed using JMP Pro software. Significance was tested using ANOVA and Tukey’s-HSD pairwise comparisons. Leaf sugar and starch data was analyzed using JMP Pro software. Significance was tested using Student’s t test.

## SUPPLEMENTAL MATERIAL

**Figure S1.** Diagram of leaf disc sampling and grafting experiment.

**Figure S2.** Fruit BRIX and yield of *bip* mutants.

**Figure S3.** Normalized expression of *GPT2* in M82 and *SiFT*.

**Figure S4.** Promoter sequence alignment of *GPT2*.

**Figure S5.** Diffcorr networks for bipvs.Lu and HBvs.LB.

**Figure S6.** Gene expression patterns of Diffcorr genes.

**Table S1.** Differentially expressed genes for *bip0663* vs. Lukullus and HB vs. LB.

**Table S2.** GO analysis of 57 shared DEGs.

**Table S3.** Literature-curated vascular related genes.

**Table S4.** 11 shared Diffcorr genes.

**Table S5.** GO terms for HB vs. LB DEGs.

**Table S6.** GO terms for genes highly correlated with *GPT2*.

**Table S7.** GO analysis of 5 gene co-expression network communities.

## DATA DEPOSITION

RNA-seq and WGS libraries are deposited in NCBI’s SRA database (Accession number PRJNA703300). R scripts used for data analysis and network construction can be found on GitHub (https://github.com/karoczar/Sinha-Lab-Scripts).

## ACKNOWLEDGMENTS

We acknowledge help from Mary Lee, Kirsten Brand, Gabriel Luis Moreira, Kristina Khuu, Eduardo Ramirez, Divya Kumaria, and Amber M. Flores in field sample, leaf and BRIX data collection, and seed collection. We also thank John Harada, Leonardo Jo, Rie Uzawa, Siyu Li, and members of the Sinha Lab for advice. We also thank CyVerse for online data storage. Support from USDA-NIFA (grant no. 2014-67013-21700) to NRS, Rosalinde H. Russell Fellowship to SDR, and JSPS KAKENHI (19K23742, 20K06682) and a JSPS Fellowship (13J00161) to HN is gratefully acknowledged.

